# Slow oscillations provide the spatio-temporal framework for long-range neural communication during sleep

**DOI:** 10.1101/2021.10.03.462961

**Authors:** Hamid Niknazar, Paola Malerba, Sara C. Mednick

## Abstract

Slow oscillations (SOs, <1Hz) during non-rapid eye movement sleep are thought to reflect sleep homeostasis and support memory consolidation. Yet, the fundamental properties of SOs and their impact on neural network communication are not understood. We used effective connectivity to estimate causal information flow across the electrode manifold during SOs and found two peak of information flow in specific phases of the SO. We show causal communication during non-rapid eye movement sleep peaks during specific phases of the SO, but only across long distances. We confirmed this prediction by cluster analysis demonstrating greater flow in global, compared with local, SOs. Finally, we tested the functional significance of these results by examining which SO properties supported overnight episodic memory improvement, with the underlying assumption that memory consolidation would engage global, long-range communication. Indeed, episodic memory improvement was predicted only by the SO properties with greatest causal information flow, i.e., longest distances between sinks and sources and global, but not local, SOs. These findings explain how NREM sleep (characterized as a state of low brain connectivity) leverages SO-induced selective information flow to coordinate a wide network of brain regions during memory formation.

## Introduction

Human brain oscillations measured by electroencephalography (EEG) reflect synchronized activity of thousands to millions of neurons, with spike timing modulated by the phase of ongoing oscillations ^1^. Current models propose that oscillations, separately or in coordination with one another, serve to organize information processing and communication in neuronal cortical networks in a state-dependent manner ^2^ and are thought to be markers of neural processing that occur in relation to specific behavior. A growing body of studies using neuronal recordings ^3^, cortical and local field potential ^4^, EEG signals, cognitive and behavioral methods ^5^ have demonstrated the important and causal role of non-rapid eye movement (NREM) slow oscillations (SOs, <1Hz) in brain function. While SOs can be found in small cortical networks [decorticated cats papers of Timofeev], naturally occurring SOs are considered a global brain network phenomenon, engaging neurons throughout the cortex, as well as several subcortical areas, including the thalamus, striatum, and the cerebellum ^6^.

Studies typically analyze SOs using functional connectivity to measure temporal similarity or correlations between different EEG channels ^7^, which is limited to correlational measures, and cannot identify directional causal communication. In contrast, effective connectivity tests the influence that one neural system exerts over another either directly or indirectly ^8^ estimated using Granger causality. According to this approach, a causal relation is detected if past values of a primary signal (source) help predict a second signal (sink) beyond the information contained in its past alone ^9^. Granger causality and causal information flow can be quantified using a multivariate vector autoregressive model (MVAR) and tested using the coefficients of the fitted model^9^. The current study adopted the generalized form of partial directed coherence ^10^ (GPDC) ^11^ to quantify causal information flow during the SO across EEG channels. We specified two quantifiers: outflow (from a source to other EEG channels) and flow (between sources and sinks), and how distances between SO origin channels, sources, and sinks modulated the information flow.

Slow oscillations are generally understood as large travelling waves that coordinate activity across cortical regions and in cortical-subcortical interactions ^12,13^ and have been shown to support memory formation^14,15^. One potential way they accomplish this process is by changing communication networks via synchronized neuronal activity along efferent pathways spread across a broad range of brain areas relevant to memory consolidation and memory replay ^16^. Still, the fundamental properties of SOs that facilitate changes to communication networks and support memory consolidation remain unexamined. The current study investigated the SO properties that reflect changes in the information flow across the brain and their impact on memory formation.

### Causal information outflow from sources during SOs

We used generalized partial directed coherence (GPDC) ^11^ as an estimator of causal information flow between different EEG channels. Four EEG channels (Fz, Cz, Pz, POz) were considered as the source of information flow and 12 channels (three channels in each region of frontal, central, parietal and occipital), were considered as the sink of information flow. An automatic system detected SOs in the source channels and two quantifiers were computed in the window 1sec before to 1sec after the SOs trough (Figure 1.A). We defined the first quantifier as the outflow from a source (*CH*_*outflow*_, ‘channel outflow’) and the second quantifier as flow from a source channel to a sink region (*CH* → *R*). Both quantifiers were estimated with respect to the SO phase and computed as an average for each subject (see Methods for the definitions and details). Figure 1.B shows the average and standard deviation of *CH*_*outflow*_ for each source across SO channels and based on SO phase. As SO phase changes, outflow peaks at − *π* / 2, drops to a local minimum at 0 (corresponding to the SO trough) and peaks again at *π* / 2. As a comparison, we also estimated outflow from sources in non-SO windows with the same number of SOs in each stage. Figure 1.C displays the outflow values for each source in pre- (phase = − *π* / 2) and post-peaks (phase = *π* / 2), the SO trough (phase = 0), and non-SO windows. We tested outflow differences in SO and non-SO windows with ANOVA and post-hoc analysis. The results showed that outflow (*CH*_*outflow*_) in the pre- and post-peaks was significantly larger than outflow in non-SO windows (p-value<0.001), but outflows at SO troughs and the non-SO windows were not different (p-value>0.05) (Figure 1.C). Also, compared to non-SO windows, there was no significant difference between outflow in half a cycle after SO trough (phase = *π*) and significant or marginally significant (0.05<p-value<0.1) difference between outflow in - *π* phase, which can be caused by the effect of a prior SO on the successive SO in a sequence of SOs.

**Figure 1.**
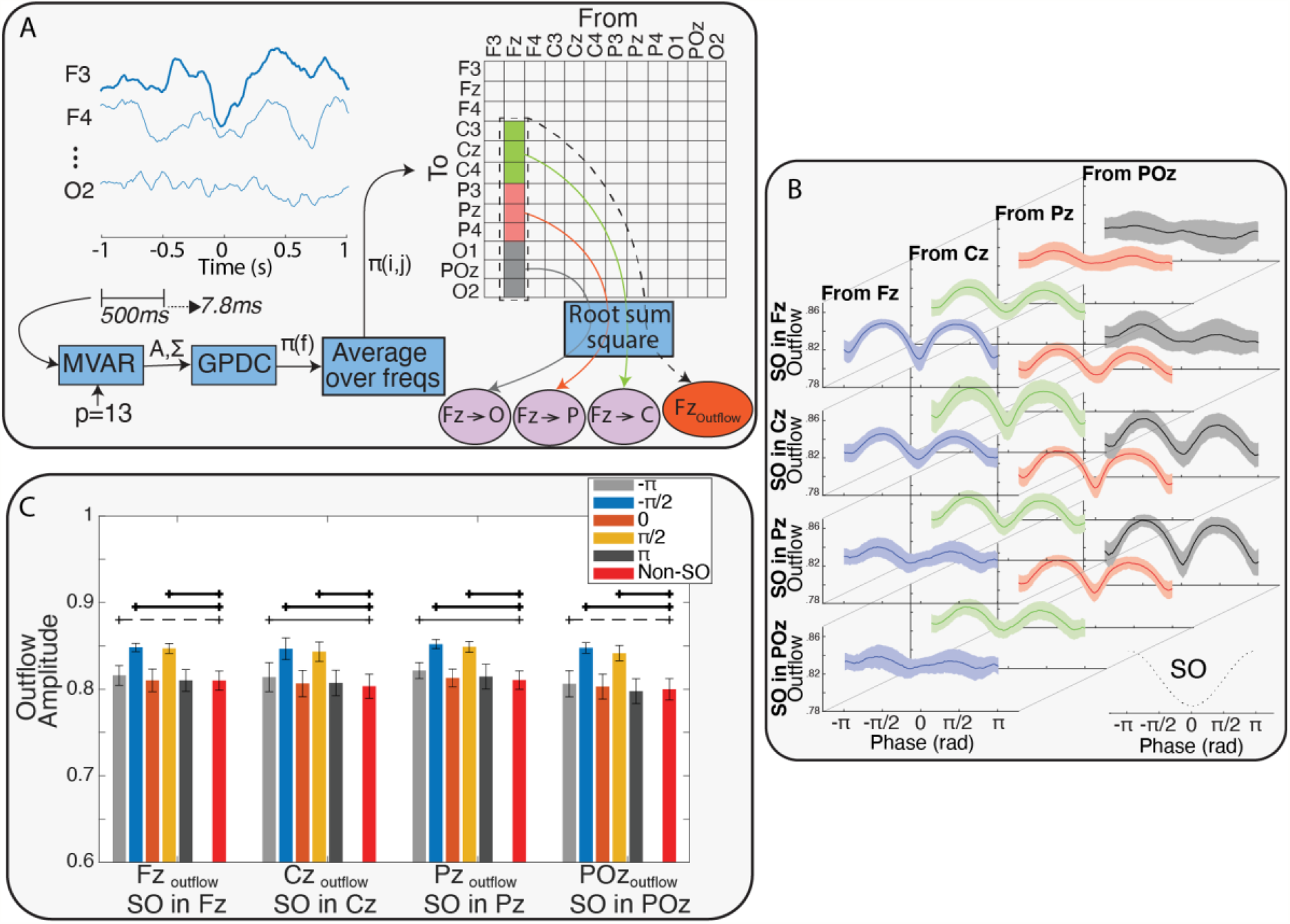
A. A representation of the process that estimates causal information flow between EEG channels. In summary, after detecting SOs, a multivariate autoregressive model was fit on each 500ms window of 12 EEG channels from 1sec before to 1sec after each SO trough, with stride of 7.8ms. By using the GPDC method and averaging over all frequency, the effective connectivity matrix (*π*) and the two defined quantifiers were calculated. After resampling the quantifiers time series based on the SO phase, the phase series were averaged over all SOs of each subject to obtain the phase series of 1) the outflow from the sources (e.g. Fz_Outflow_) and 2) the flow from sources to sinks (e.g. Fz→Cz) for each subject. B. Variation of outflow from all sources conditioned on the SO channel. Each row shows the outflow variation across the subjects over phase of SOs when the SO was detected in the represented channel. In Each row, the columns show outflow variation in each of the sources (mean±std). The phase series and their variation across the sources and SO channel show peaks before and after the SO trough and a possible relation between source and SO channel. The results in this figure suggest that sources closer to SO channels had greater height in outflow peaks. C. Comparing the outflows at five different phases of the SOs (SO phase={−π, π / 2, 0, −π / 2, π}) to outflow in non-SO windows. ANOVA and post-hoc tests found that peaks in phase of − π / 2 (pre-peak) and π / 2 (post-peak) had significantly larger outflow, while there were no significant differences between outflow in non-SO windows and SO trough. Thick black lines indicate a strong significant difference (p-value<.005), thin lines indicate significant difference (p-value<0.05) and dash lines indicate marginal difference (0.05<p-value<0.1) between SO and non-SO windows.

In a linear mixed-effects (LME) model, we examined the effect of sources, SO channels, and distances between sources and SO channels 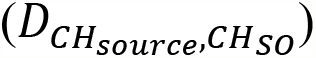 and peak phase (i.e., whether the peak preceded or followed the SO trough) on the height of the peaks in information outflow. We assigned discrete values for the distances 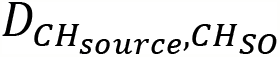 between source (Fz, Cz, Pz and POz)and SO channel as zero to three in the linear model, adding one for each ‘step’ in the frontal-occipital axis. We used a simple model to calculate the distances as we considered distance between each two neighbor regions equal to 1 (This meant that if *CH*_*source*_ was Fz and *CH*_*SO*_ was Cz, difference would be 1, and if *CH*_*source*_ was Cz and *CH*_*SO*_ was POz, the distance would be 2). Table S.1 presents the coefficients and related p-value identified by the LME model.

Based on LME modeling outcome, there were significant linear effects of the SO channel (p-value<0.05), distance between the SO channel and the source 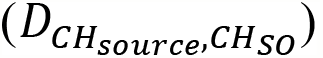, and phase (pre-/post-peak) on the height of outflow peaks (p-values<0.05), with 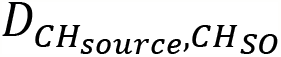 having the largest linear effects (higher coefficients). The negative coefficient of 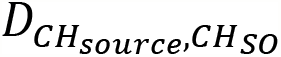 demonstrated that information outflow increased with proximity of the sources to the SO channel. For example, for an SO detected at the Fz channel, the lowest peak of causal information outflow would be found in the POz channel, whereas moving closer to the Fz channel, the peak of the outflow would increase, with highest outflow at Fz itself. The positive coefficient of the SO channel indicates that the position of the SO (anterior vs posterior) modulated the height of the outflow, where SOs in the anterior channels may produce lower outflow in all sources compared with SOs in more posterior regions. Additionally, the negative coefficient of peak phase (i.e. being a pre- or a post-peak) indicates that pre-peaks are expected to have greater outflow than post-peaks.

### Causal information flow from sources to sinks during SOs

In the previous section we showed that the SO-related information outflow from a source was significantly affected by the distance between the SO channel and the source, where sources closer to the SO had largest peaks. To analyze how information flow depended on sender and receiver, we next focused on source – sink pairs. Causal information flow from each source to each sink (*CH* → *R* quantifier, where *Fz* → *P* would have *Fz* as source and parietal as sink) were computed over all SOs. For each SO channel, we averaged all *CH* → *R* phase series in each source–sink pair within each subject. Figure 2.A presents samples of *CH* → *R* phase series variation from two of the sources (Fz and POz) to three sinks separately (central, parietal and occipital with Fz source and frontal, central and parietal with POz source), all cases with SO channel in Fz (see Figure S.2 to S.5 for comprehensive results).

**Figure 2.**
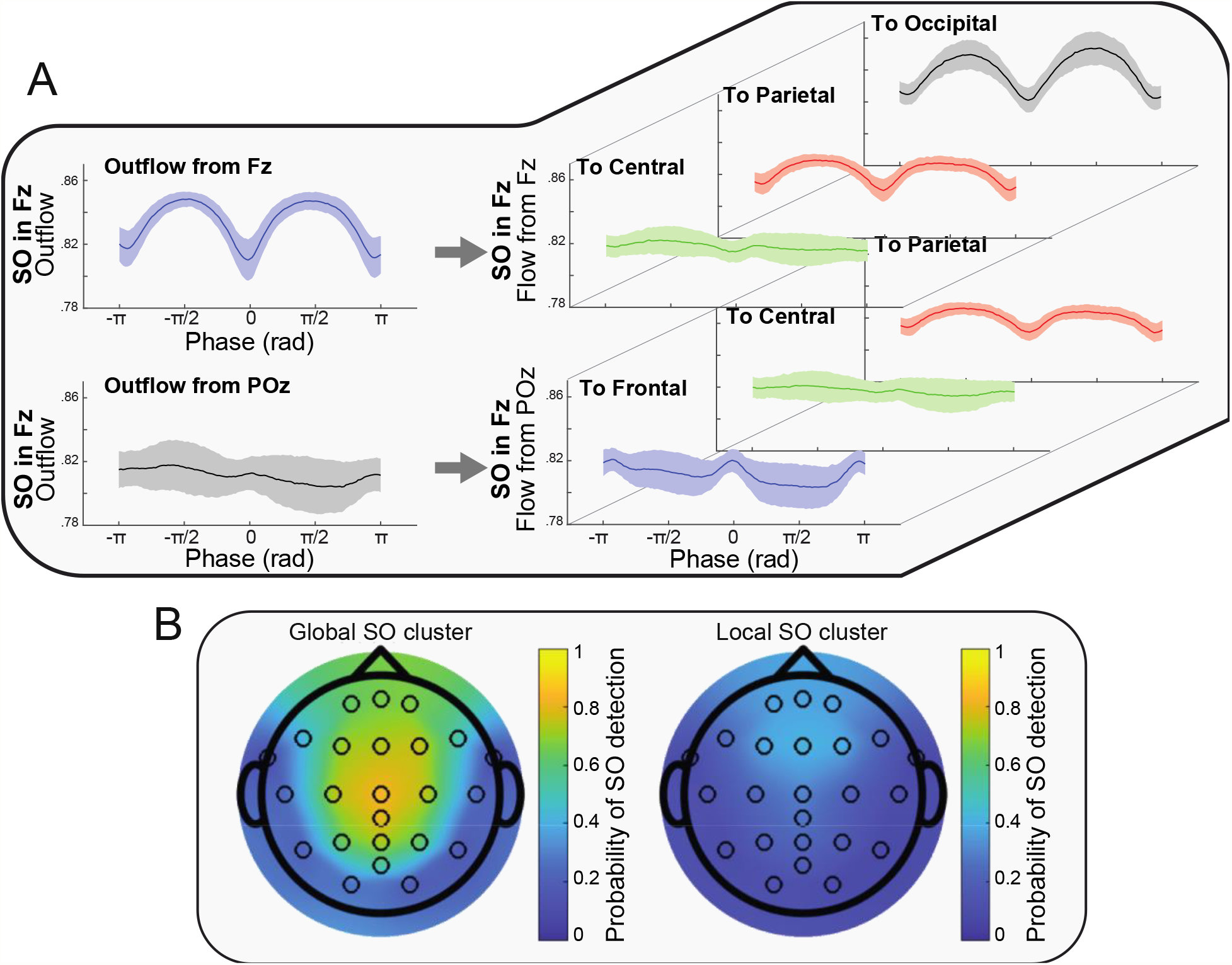
A. Examples of information flow from source to sink (CH → R quantifier). The left plots show the total outflow from the specified source and with presence of SO in the specified channel (Samples from Figure 1.B). The right plots represent the portions of the flow from the source to the different sinks. For example, the top-left plot shows the outflow from the Fz as the source when there were SOs in Fz channel and the three top-right plots represent portions of that flow to each of the sinks. By considering the topology of the SO channel, source and sinks the results suggest a relation between distance of sink to SO channel and the peaks of information flows. B. Two clusters of SOs found by the clustering method (Cluster 1: Global SOs, Cluster 2: Local SOs). The colors represent the density of the SOs in each channel.

Observing the *CH* → *R* quantifier values in Figure 2.A (and Figure S.2 to S.5), we noticed that there was a relation between distance of the sink to SO channel 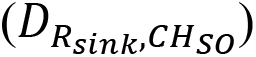 and height of the flow peaks. This suggested that the causal information flow (*CH* → *R*) could be modulated by which source–sink pair was considered, the distance of source–sink pairs to the SO channel, and the distance between the source and the sink in each pair. To investigate these potential effects, an LME model was fitted to the *CH* → *R* values in the peaks of the flow (pre- and post-SO trough), with linear predictors: SO channel, source–sink pairs, distance of source–sink pairs to the SO channel 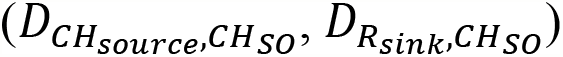 and distance between source–sink pairs 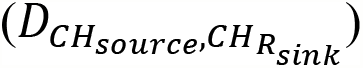.

The outcome of our LME model (Table S.2) demonstrated a significant linear effect of 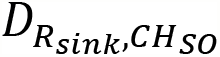 and 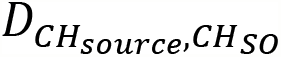 on the height of the flow peaks. Also, there was significant effect of peak phase (pre- or post-trough) where the height of the peak in the pre-trough phase was higher than in the post-trough phase (coefficient<0). In contrast, we found no statistically significant linear effects of SO channel, source, or distance between source–sink pairs on the height of the causal information flow. The coefficients values showed different magnitude and direction of the effect in each predictor. The positive coefficient of distance between sink and SO channel and negative coefficient of distance between source and SO channel showed closer source to SO channel (less distance) and farther sink to SO channel (larger distance) had higher peak of information flow in comparison to other compositions. For example, an SO in Fz channel, sources closest to the SO channel and sinks farthest from the SO channel (e.g., *Fz* → *O*) have the highest information flow, suggesting that brain communication during the SO is highest at locations far from the SO, and distance between SO channel to the source–sink pair impacts causal flow.

Based on outcomes from our analysis, the following points can be inferred:

- Sources closer to the SO channel send the greatest magnitude of information (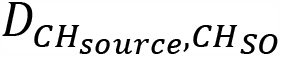 coefficient for flow peaks <0).
- Sinks farther from the SO channel receive the greatest magnitude of information (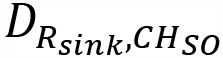 coefficient for flow peaks > 0).
- The distance between the sink and the SO channel shows strongest effect on information flow compared with all other predictors (absolute value of 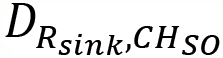 coefficient for flow peaks >> absolute value of other predictor coefficients).

### Effect of SO clusters on causal information flow

So far, we have shown that the impact SOs on causal information flow depends on the distance between the source to the SO channel when considering outflow, and depends on the distance of the sink to the SO channel when looking at specific flow between source-sink pairs. Also, channels closer to the SO channel can send more causal information flow, whereas channels farther from the SO channel can receive more causal information flow. In light of these properties, we hypothesized that the primary benefit of SO’s effect on brain information processing is to permit causal communication between remote brain regions. If this was true, it follows logically that global SOs (SOs that propagate across a large portion of the scalp) should mediate more information flow compared to other types of SOs. Studies investigating the spatial and temporal co-occurrences of SOs report that the majority of SOs are traveling waves, with their pattern of origin and propagation providing an outline of cortical connectivity ^12^. Using a cluster-based analysis of SO co-occurrences across the EEG channel manifold, prior work from our group identified three spatio-temporal categories: frontal, local and global ^13^, whereby global SOs occur in most electrodes, local SOs occur in few electrodes without location specificity, and frontal SOs are confined to the frontal region. Along with having greater footprint and larger amplitude, global SOs also provided greater hierarchical nesting of thalamocortical sleep spindles, suggesting functional differences between SO categories, with global SOs more poised to activate systems consolidation ^13^.

We tested the hypothesis that global SOs should have greater causal flow than non-global (local) SOs by comparing the effective connectivity in each type. We first clustered global and non-global SOs using a method previously introduced by our team ^13^. Figure 2.B shows the occurrence rate of SO in each of the channels and in each of the two clusters on the scalp surface after the clustering process. Based on results from ^13^ and our presented results in Figure 2.B, the two detected clusters were interpreted as global and local SOs clusters. To test if there was an effect of cluster identity (i.e. being a global or local SO) on the height of information flow peaks, we added cluster identity (global as 1 and local as 2) as a fixed effect along with other used predictors in a LME model of information flow peak height.

The results of LME models for peaks of information flow from the sources to the sinks (Table S.3) showed the same effects of the previous predictors but also a significant linear effect of clusters on the height of the flow peaks (p-values < 0.05). Negative coefficients for the cluster effect showed that peak heights were higher in global SOs (cluster = 1) than local SOs (cluster = 2). We interpret this as confirming our hypothesis that global SOs mediate larger information flow compared to other SO types.

### Long-term memory improvement and causal information flow during SOs

Next, we investigated the consequence of the SO properties on the formation of long-term memories. Based on the importance of down-to-up state transitions ^17–21^, matched to the post-peak in our study, we examined the relation between causal information flow and word-pair association (WPA) improvement during the post-peak (peak of information flow at *π* / 2 after the SOs trough). WPA improvement was measured as the ratio of performance after the sleep night (post-sleep) to the performance before the sleep night (pre-sleep). For each subject, we assigned the post-peak outflows to one of the four distances between the source and the SO channel 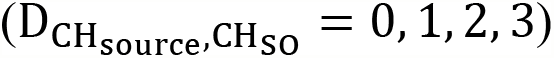 and then averaged within each group. We then evaluated linear relations between the outflow in each group and WPA improvement using linear regression models (the p-values adjusted to Bonferroni correction) (Figure 3).

**Figure 3.**
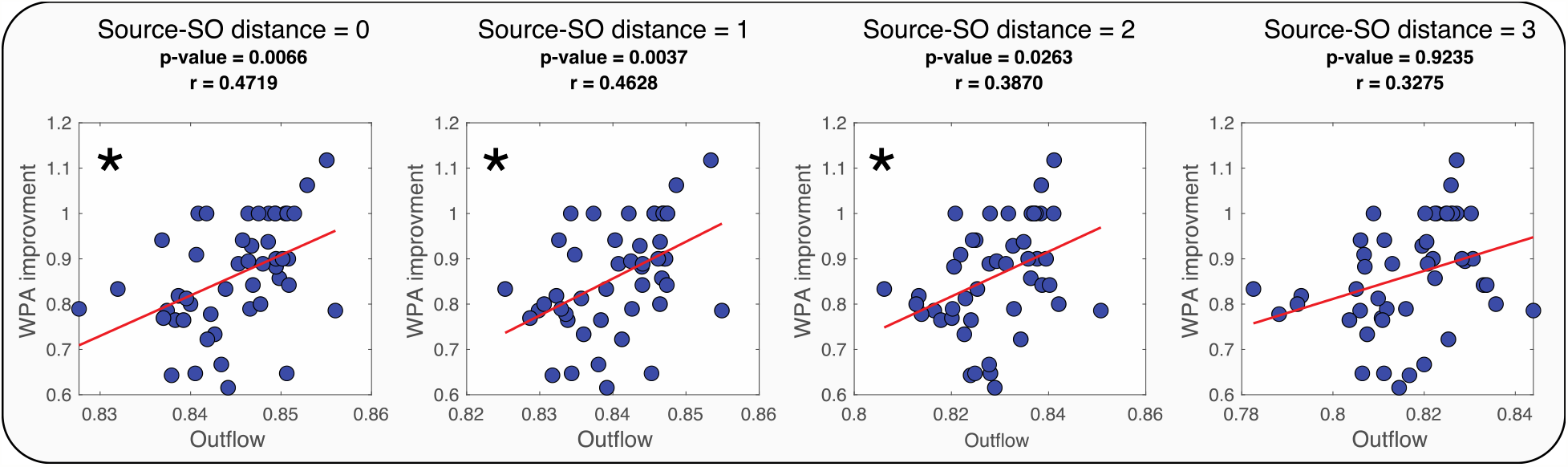
Regression test for outflow of source and WPA improvement in four different conditions of distance between SO channel and outflow source 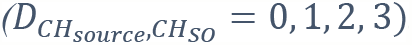. In each, we show the p-value, the linear coefficient is reported as r and the significant linear relationships are marked with asterisks.

Results in Figure 3 show that memory improvement was associated with the distance between SO and sources, with a significant positive linear relation between WPA improvement and outflow when the source is close to the SO channel 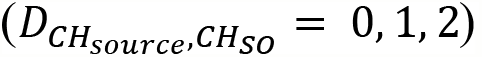, but not when the source is far from SO channel 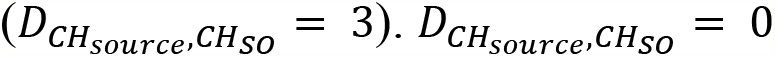 had the greatest r-value in the regression test and the r-value got smaller as distance between the channels increased 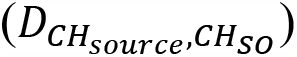: this can be interpreted as the outflow from sources close to SO was a better predictor of WPA improvement.

Next, we examined how the distance between source and sink channels, and the relative distance between the source and sink to the SO channel impacted WPA improvement (Figure 4.A). First, we defined three possible distances between sink and source 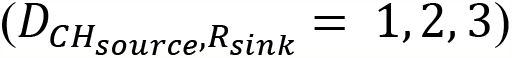 and averaged flows from sources to sinks which had the same distance. Figure 4.B displays the linear relation tests of WPA improvement and averaged flows in each specified distance between sink and source.

**Figure 4.**
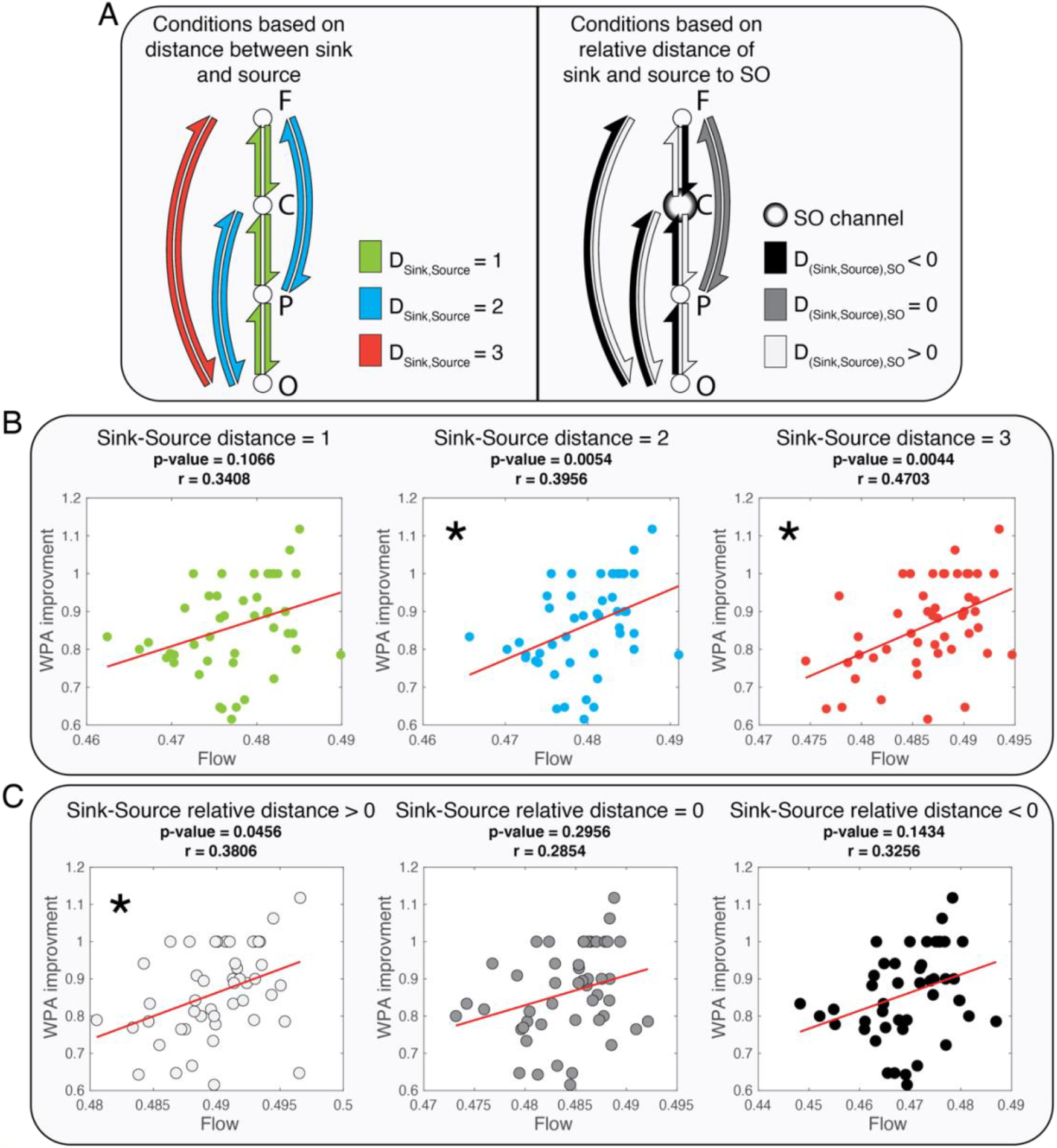
Linear relationships between flow and WPA improvement based on different conditions of distance between sink and source, and relative distance of sink and source to SO channel. The asterisks mark significant linear relationships. A. A representation of the three condition of distance between sink and source (left graph) and an example of three possible conditions of relative distance of source and sink to SO channel (right graph, for SO channel at Cz). In the left graph each color represents pairs of sink and source of information flow with the same distances. The right graph shows an example of the conditions of the relative distance when the SO channel is Cz. The relative distance is greater than 0 when source is closer to SO channel than sink to SO channel and smaller than zero when sink is closer to SO channel than source to SO channel. B Results of correlation and regression test of relation between the flow and WPA improvement for three conditions of distance between source and sink 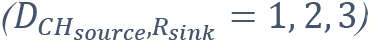. C. Results of correlation and regression test of relation between the flow and WPA improvement for three conditions of relative distance between source and sink to SO channel.

Based on the results presented in Figure 4.B, larger distances between sinks and sources 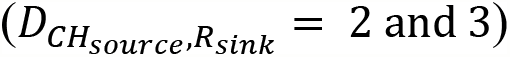 were linked to significant positive linear relation between information flow and WPA improvement. The r-value increased with greater distance between sink and source suggesting a stronger linear relation between information flow and WPA improvement for long-range communications.

Next, we examined the association of WPA improvement and flow in different relative distances from the sink and source to SO channel. We defined the relative distances as the difference of distance between sink and SO channel and distance between source and SO channel 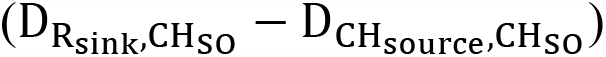. Hence, relative distances greater than zero indicated that the source was closer than the sink to the SO channel. For example, if Cz was the SO channel, the relative distance of Fz source and occipital sink would be greater than zero, as 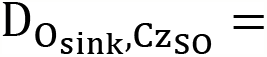 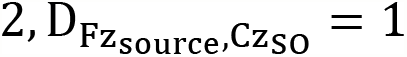. Figure 4.C shows a significant linear relation between WPA improvement and causal information flow only when the relative distance is greater than zero. Together, these results are consistent with our proposed hypothesis that long-range communication mediated by SOs plays an important role in memory consolidation.

A final test of this hypothesis investigated whether global, but not local, SOs mediated the relation between causal information flow in SOs and WPA improvement. We first measured outflow in the post-peak SO phase within global and local SO clusters and calculated significance levels of regressions between outflows and WPA improvement, shown in Figure 5.C (refer to supplementary for the detailed results, Figure S.6 and S.7). We found a greater number of channels with a significant linear relation between their outflow and WPA improvement in the global cluster compared with the local cluster (4 combinations of source and SO channel in the local cluster and 13 combinations of source and SO channel in the global cluster, Figure 5.C).

**Figure 5.**
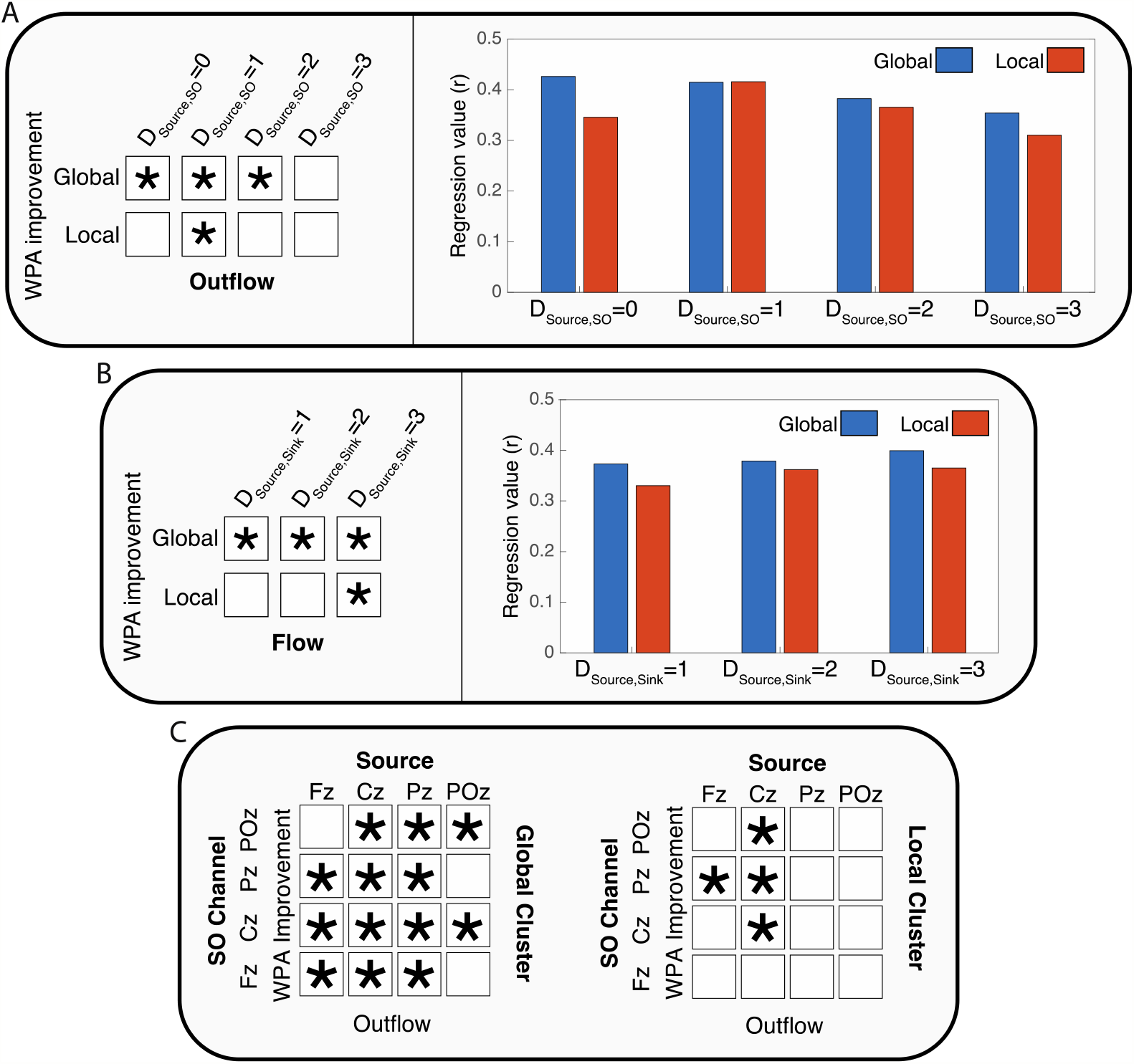
The effect SO clusters (local and global) on the relation between causal information flow and WPA improvement. The asterisk shows significant linear relationships. A. Results of correlation and regression tests for four distances between source and SO channels 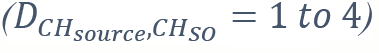. The left panel present the conditions that there was significant linear relation between outflow and WPA improvement. The right panel shows the r-values of regression in different conditions of distance of source and SO channel in the two clusters. B. Results of correlation and regression tests for three distances between sources and sinks 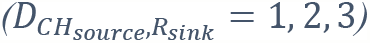. C. Summarized results of the linear relations between outflow from source and SO channel combinations for global and local SO clusters. Detailed results are presented in Figure S.5 and S.6.

Next, we averaged the outflows based on the distances between sources and SO channels in each cluster and examined correlations with WPA improvement. Given our prior result that short distances between source and SO channels have the highest information flow, we expected that global clusters with shorter distances between these channels would show the strongest associations between outflow and memory improvement. Indeed, our results showed significant relations between outflow in the global cluster when the distances between SO and sources are smaller than 4. In the local cluster, consistent with the reduced footprint of SOs in this cluster, we found a significant relation between outflow and WPA improvement only when the distance between SO and source was equal to 1 (Figure 5.A). Furthermore, r-value comparisons showed that the best condition (greatest r-value) for modeling WPA improvement was in the smallest distance between SO and source and for SOs in the global cluster.

We then conducted the same analysis between sink and source distances in each cluster. In each of the clusters, we calculated and averaged flows post-peak across distances between sink and source 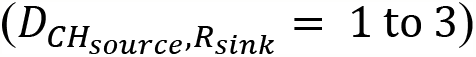. We found (Figure 5.B) a significant linear relation between flow and WPA improvement for all sink/source distances in the global cluster. Conversely, in the local cluster there was a significant relation between flow and WPA improvement only when sink and source were at the farthest distance 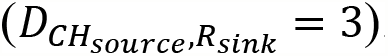. In sum, these results demonstrate that the best condition for modeling WPA improvement based on information flow is during global SOs across long-range neural channels.

## Discussion

In this work, we investigated temporal and topographical properties of the sleep slow oscillation (SO) using effective connectivity, which establishes directional causality of information flow. We estimated information outflow from individual channels and flow between channels. We found two peaks in outflow before and after the SO trough, while magnitude of outflow at the SO trough was similar to outflow in non-SO windows. By considering source and sink quantifiers, we determined that sources closer to the origin of the SO were the biggest senders of information, specifically during the pre- and post-peaks. Additionally, sinks farthest from the SO channel were the highest receivers. Global SOs also mediated larger information flow compared to local SOs. Taken together, our effective connectivity findings suggest that one fundamental property of the SO is to facilitate long-distance causal communications across brain areas. Next, we tested the functional implications of this property by examining the temporal and topographic conditions that predict overnight memory improvement. Our results show that conditions that minimize the distance between source and SO channels and maximize the distance between sink and source channels, as well as engage global SOs, are most strongly associated with memory improvement. Our findings are consistent with the notion that SO-driven long-range communication supports long-term memory formation.

### SOs shape the causal communication network in NREM sleep

SO waveform peaks have been linked to neural firing rate and spiking activity via multi- and single-unit recording and intracranial EEG ^22^. Nir et al. (2011) showed the mean firing rate of units in some brain regions (e.g., orbitofrontal cortex, anterior cingulate, supplementary motor, parietal cortex, parahippocampal gyrus, and hippocampus) increased up to 200% 500ms before and after the SO trough, and decreased down to 40% within the SO trough in comparison with the mean firing rate in NREM (N2 and SWS). Although higher firing rate and spiking activity has been interpreted to signify greater potential outflow of information ^23^, the precise nature of the SO-triggered information flow has yet to be identified. In the current work, we found two peaks of information outflow (pre- and post-SO trough), whereas causal information outflow at the SO trough showed the same magnitude as that of the non-SO windows. However, the timing and phase of the outflow peaks in our study (± *π* / 2 and about ±250ms from the SO trough) did not match the peak firing rate reported by Nir et al. (2011) (±*π* and about ±500ms from the SO trough), suggesting that there are other components and conditions which affect causal information flow in addition to spiking activity.

Global brain effective connectivity and information flow are modulated by the transition from wake to sleep. Using transcranial magnetic stimulation (TMS) to probe propagation patterns across wake and sleep stages, Massimini et al. showed the lowest levels of brain connectivity during NREM sleep ^24^. Furthermore, effective connectivity analysis using functional magnetic resonance imaging (fMRI) and simultaneous EEG recordings have described N2 as an unstable network and SWS as a stable network ^25^. Together with our current findings of equivalently low levels of information outflow during SO troughs and non-SO windows during NREM, one emergent hypothesis is that causal communication across brain regions during NREM sleep is overall poor, and punctuated by bursts of selective high-communication events, mediated by SOs.

### Causal information outflow depends on distance from SO channel

In the previous section we discussed the dependency of causal information outflow on the time and phase of the SO. Our LME modeling results support the possibility that SOs enable information flow at selected times and between selected locations. In particular, the distance between source and SO channels had the strongest effect on peak height (i.e. amount of information flow). Functional connectivity approaches have characterized SOs as a local phenomenon, measured in their direct effect on small neural regions or less than 20% of all EEG channels ^26^. SOs are also associated with localized brain activity, suggesting that brain areas near the SO channel have a higher potential to send information (i.e., higher firing rate). Our results further suggest that the closer the source, or information sender, is to the SO channel, the higher the information outflow.

Our findings also suggest differences between the pre- and post-SO trough peaks. One potential source for these differences may be that we combined SOs of different types, including single and sequential SOs ^27^, wherein sequential SOs have pre- and post-SO peaks that are temporally linked with each other. In these cases, the pre-peaks may be influenced by the preceding SO and the current SO, whereas post-trough peaks would be only affected by the current SO. Additionally, studies have shown functional differences between pre- and post-SO activity, including greater synchronization between SOs and spindles ^28,29^ and more efficient cueing during targeted memory reactivation (TMR) during the post-SO trough peak ^21^. Thus, our data support the notion that there are functional and effective connectivity differences between SO peaks, but that further research is needed to tease apart properties of single or sequential SOs.

By modeling the outflow peaks, we showed a significant effect of the SO channel, with the positive coefficient demonstrating an increase in the height of outflow from anterior to posterior regions. These results are likely due to the presence of significantly fewer SO in posterior regions, suggesting that total information flow is not higher in the posterior channel in general, but rather that during the few posterior SOs that do occur, there is a higher amount of information outflow than other areas. Future research may determine why outflow increases as the number of detected SOs diminish.

### SOs facilitate causal communication between more distant regions

The SO has also been described as a global phenomenon that recruits the entire cortical network ^30^. We, therefore, probed information flow by considering source and sink channels independently and together as quantifiers and discovered that regions farther from the SO channel are more engaged in receiving information. This suggests that the global brain communication during the SO may be organized with local areas near the SO sending more information and local areas farther from the SO receiving more information. As such, brain areas engaged in high causal information outflow (i.e., sending information) have a low chance of simultaneously receiving information. Given these results, we hypothesized that the effect of distance between sinks and sources and greater causal communication between farther regions should be specifically accentuated in the case of global SOs which involve a greater number of brain regions ^13^. We tested this hypothesis by using global or local SOs as predictors in our model and demonstrated a significant effect of SO cluster on the height of the flow peaks, with global SOs having higher peak heights than local SOs.

### SOs as a facilitator for long-term memory

One leading hypothesis of memory consolidation suggests that encoding networks get reactivated during sleep ^31^, specifically during NREM sleep^32^. Given the near-infinite diversity of the encoding network, memory reactivation requires both short and long-range information transfer across a wide range of brain areas. However NREM is a sleep period generally characterized as showing a breakdown in connectivity with increasing depth ^24^, which would not be conducive to reactivation. Our results may pose a potential solution to this seeming paradox, with SOs providing the temporal-topographical event framework whereby long-range information flow increases dramatically compared to the relatively local activity patterns of the rest of the sleep period.

It is well established that SOs provide temporal coordination for memory-related brain activity during systems consolidation ^32^ and a greater effect of TMR has been shown when cues are coupled with the down-to-up phase of the SO ^18,21^, similar to the post-peak in the current results. Given the notion of the SO as a traveling wave ^12^ that enhances long-distance causal information flow, with SO location as the mediator of this effect, our data suggest a potential mechanism for causal communication between brain regions during the reactivation process. As memories get reactivated in cortical-hippocampal networks, we predicted that the temporal and topographic SO properties elucidated from the analysis would be associated with overnight memory improvement. Indeed, our results showed significant associations between SO-driven, long-range communication networks and overnight improvement in episodic memory. Furthermore, global, rather than local, SOs mediated the relation between causal information flow and episodic memory improvement, even for sources far from the SO origin and close sink and source pairs. In agreement with this finding, for local SOs, the relation between causal flows and episodic improvement was limited to sources closest to the SO channel and long-distance pairs of sinks and sources.

In conclusion, we have evinced several fundamental properties of SO waveforms, including that the flow of information peaks immediately before and after the SO trough, that brain areas closest to the SO origin engage in sending information, while brain areas farthest from the SO engage in receiving. We have also shown that these properties together predict memory improvement in healthy adults. Next steps in this research direction are to test how nested oscillations promote information flow, as well as to investigate how these properties vary in clinical populations and aging humans.

